# Self-body recognition through a mirror: Easing spatial-consistency requirements for rubber hand illusion

**DOI:** 10.1101/691634

**Authors:** Hikaru Hasegawa, Shogo Okamoto, Ken Ito, Masayuki Hara, Noriaki Kanayama, Yoji Yamada

## Abstract

Typical experiments to induce the rubber hand illusion (RHI) require experimental participants to gaze at a fake hand while tactile stimuli are provided to both the fake and hidden actual hands in a synchronous manner. However, under such conditions, postural and apparent disagreement between a seen fake hand and hidden actual hand prevents illusory body ownership. Provided that humans recognize mirror images as copies of the real world in spite of their spatial uncertainties or incongruence, the sensory disagreement may be accepted in RHI settings if using a mirror to show a fake hand. The present study performed two experiments to reveal how self-body recognition of a fake hand feature via mirror affects the RHI. These experiments were conducted in an RHI environment involving voluntary hand movements to investigate not only body ownership but also agency. The first experiment (Experiment 1) examined whether illusory ownership of a fake hand seen in a mirror could be induced. Then, we examined whether the RHI using a mirror image allows disagreement in orientation between the rubber and actual hands (Experiment 2). Subjective evaluations using a RHI questionnaire demonstrated that evoked embodiment of the rubber hand was stronger in the presence of a mirror than in the absence of it (Experiment 1) and that participants experienced the RHI even if the actual and rubber hands were incongruent in terms of orientation (45 °; Experiment 2). No significant difference was found in the change of perceived finger location (proprioceptive drift) between these experiments. These findings suggest that the use of a mirror masks subtle spatial incongruency or degrades the contribution of visual cues for spatial recognition and facilitates multisensory integration for bodily illusions.

## Introduction

The rubber hand illusion (RHI) is an illusory experience of body ownership of a fake hand when the visible fake hand and occluded veridical hand are exposed to spatially and temporarily congruent visuotactile stimuli. Since Botvinick and Cohen [1] reported this phenomenon, the paradigm of the RHI has been used in studies on multisensory integration and body ownership. In this paradigm, congruency among multiple sensory cues is important [2, 3]. Smaller spatial and temporal contradictions between the actual and fake hands lead to more intense illusions of body ownership. For instance, when the fake and actual hands are spatially incongruent (e.g., when they have different postures), the body ownership illusion is weakened [4–8]. According to Constantini and Haggard, a hand angle mismatch of 20 ° between the veridical and fake hands distracts from the body ownership illusion [6]. Nevertheless, this angle value varies among studies [8, 9].

Usually in experiments on the RHI paradigm, participants are asked to gaze at the fake hand because “RHI is understood as a visual adaptation of proprioceptive position” [5]. In contrast, in the present study, they gazed at the fake hand through a mirror. Under these settings, the RHI is experienced [10–12]. Despite the reversed orientation and uncertainty of geometrical information including depth, humans recognize a mirror image as a copy of the real environment. In other words, humans accept the low reliability of the image in a mirror. This tolerance of less reliable images may also hold in RHI settings, and it is possible that incongruency between the fake and actual hands do not disturb the illusory experience of body ownership. One study supports this possibility; in it, the body ownership illusion was evoked irrespective of the egocentric or allocentric fake hand images in a mirror [11]. The present study pursues these hypothetical propositions.

In Experiment 1, we test whether the body ownership illusion is elicited when experimental participants gaze at the mirror image of a fake hand. This experiment is not merely a replication of earlier ones [10, 11] as we tested that our settings involving self-generated hand motions and tactile stimuli cause the illusory experiences. Experiment 2 investigates whether the incongruency of the fake and actual hands’ postures (angles) is accepted in the RHI setting involving a mirror. Although Kontaris and Downing [11] found that first-person and third-person perspective images of a fake hand in a mirror did not differ in terms of the subjective experience of the body-ownership illusion, they did not deal with situations where the actual and fake hands were angled. Exp. 1 and Exp. 2 were conducted in active RHI settings to investigate the body ownership and agency. Although these two senses can be distinguished during involuntary hand movements, they are strongly related during voluntary hand movements that enhance the precision of proprioceptive cues [5]. In our active RHI settings, the effect of mirror and congruency of hand posture on body ownership and agency can be investigated.

Although studies on the body ownership of phantom limbs have commonly employed a mirror (e.g., [13]), those on the RHI have thus far rarely experimented with mirrors. For instance, in the framework of the RHI, Holmes et al. used a mirror to investigate how position bias caused by a mirror affects the reaching task of an unseen hand [14]. Additionally, Preston et al. examined illusory ownership over a whole-body mannequin seen in a mirror [15]. RHI settings using a mirror may share some traits with body ownership illusions when facing an avatar image in a virtual environment [16]. In two previous studies [15, 16], whole-body ownership illusion was induced by using a mirror or images from a third-person view.

## Materials and methods

### Participants

Fifteen paid university students participated in the experiments (9 males and 6 females; mean and standard deviation of age in years: 24 and 9.2) with written informed consent. The participants were recruited using a local advertisement poster. None of the participants had prior experience of the RHI or other experiments relating to body ownership. They were unaware of the objectives of the experiments in terms of the effects of the mirror. They were allowed to take a satisfactory rest between experiments. All participants were involved in two types of experiments as described below. The experiments were conducted sequentially in the order of Exp. 1 and then Exp. 2 such that participants did not realize the expected effects of using mirror images. Hence, in Exp. 2, participants could be more familiar with the process including how they should move their hands to effectively cause the illusory experience.

### Ethics statement

This study was conducted with the approval of the internal review board, School of Engineering, Nagoya University (*#*17-12).

### Apparatus for inducing the RHI under self-generated movement

As shown in Fig 1(a, b) and Fig 2(a, b, c), the main components of the apparatus were a fake rubber hand, hand gloves, a cuboid frame, and a cloth for blindfolding that occluded each participant’s right shoulder, arm, and hand. The fake rubber hand was composed of a rubber glove containing wires and cotton. The same rubber glove was worn by participants. Two acrylic rods were fixed to the fake hand as in Fig 2(b). Participants held these rods with the thumbs and palms of right hand and moved the fake hand and their own right hands in a synchronous manner. They moved their left hands in the same manner as the fake and right hands. When the acrylic rods were removed from the fake hand, the fake hand remained still on the desktop, and an asynchronous condition was configured in which the actual and fake hands were independent. A similar setup was used in [17].

**Fig 1.**
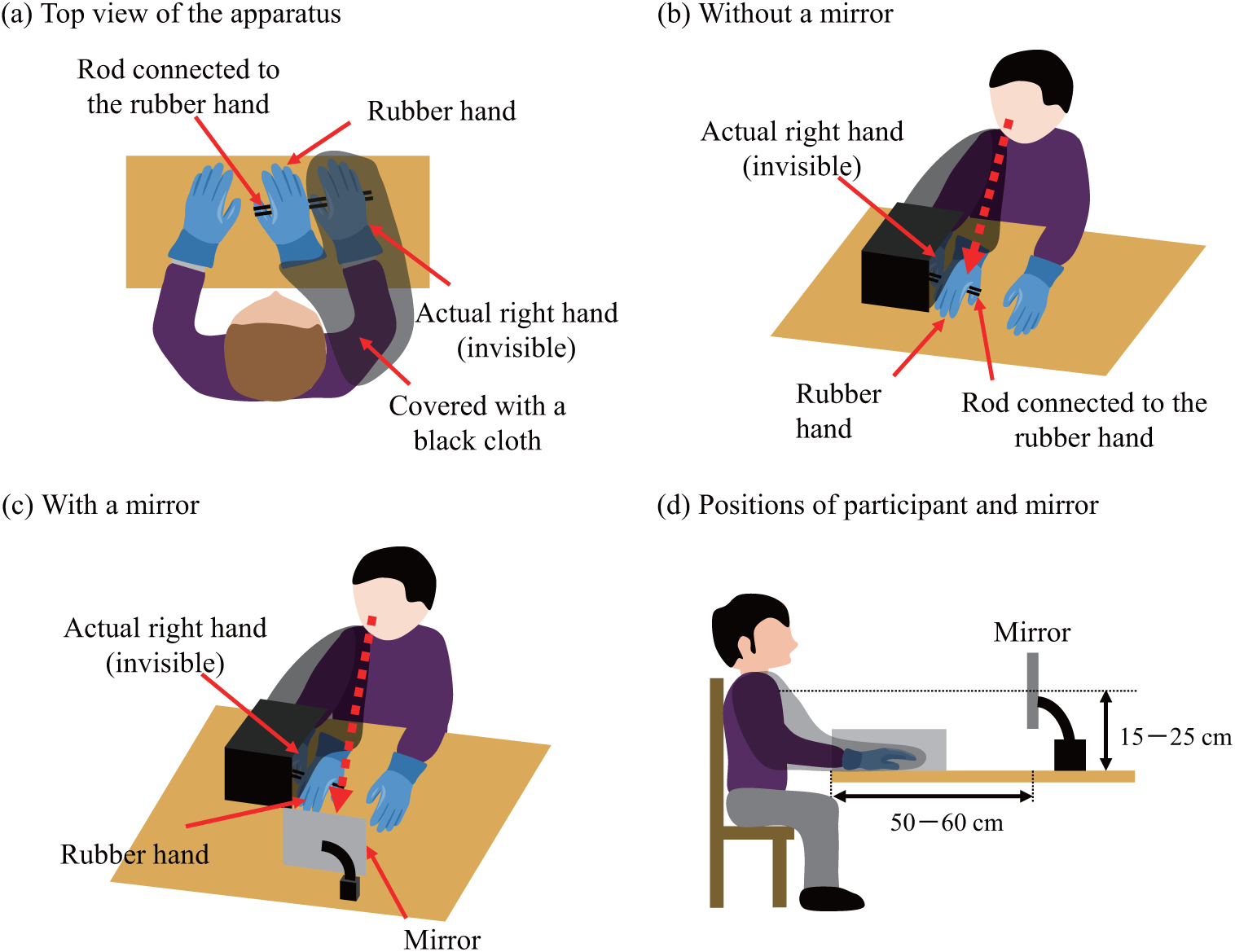
Experimental conditions. (a) Movement of rubber hand was controlled by manipulating the connected rods. (b) The actual right hand and arm were hidden by a box whereas the rubber hand was visible to participants. (c) The fake rubber hand and the actual left hand could be seen in the mirror. (d) Positions of the mirror, hand, and participant. These positions were adjusted such that participants could see their hands in the mirror.

**Fig 2.**
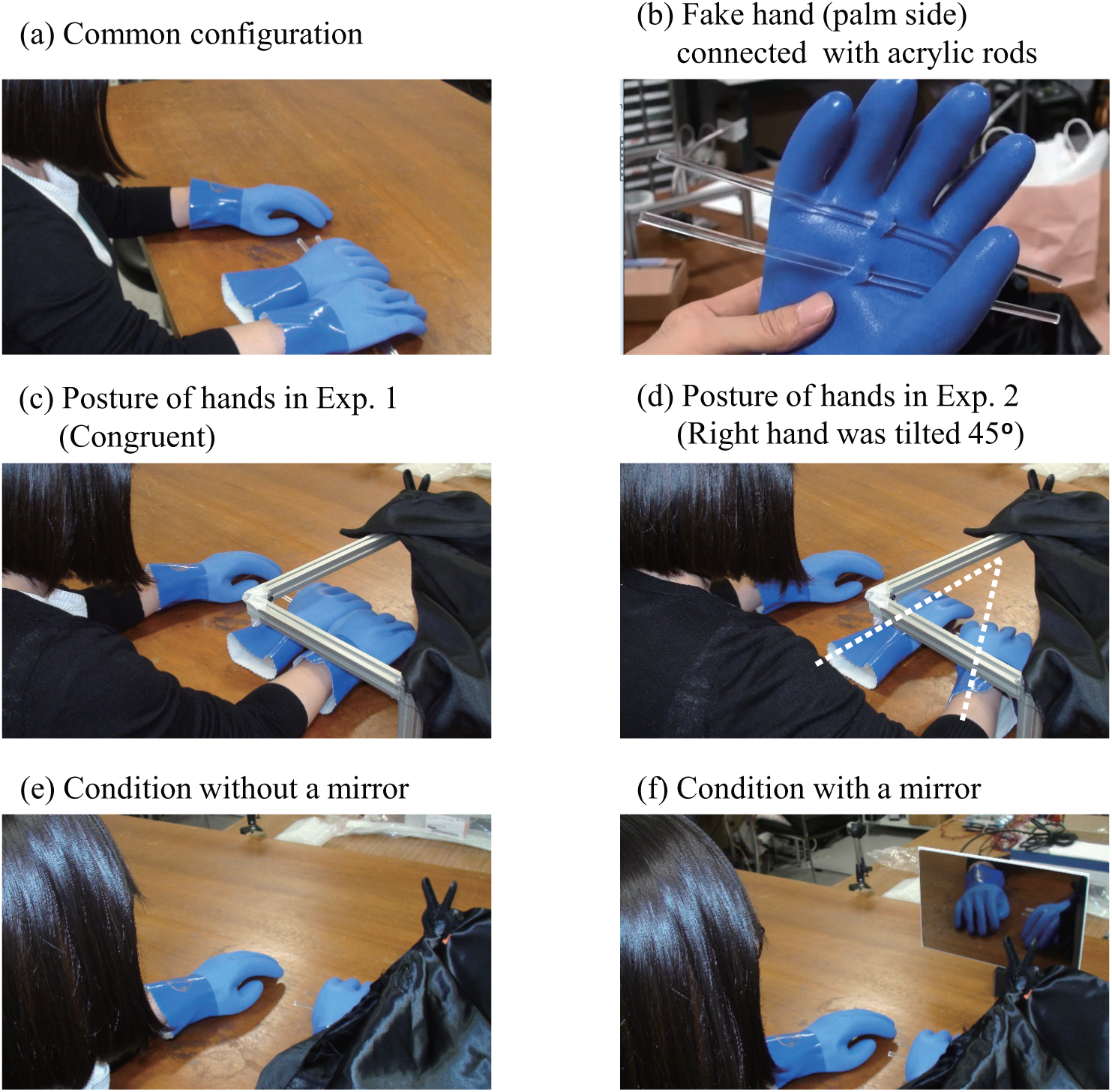
Experimental scene. (a) Common setup for all experiments. (b) Palm side of the fake hand, which was firmly supported by two acrylic rods. (c) The actual right hand was parallel with the fake hand in Experiment 1. (d) The actual right hand was angled by 45° in Experiment 2. (e) Condition without a mirror in experiments 1 and 2. (f) Condition with a mirror in experiments 1 and 2.

Fig 1(c, d) and Fig 2(f) show the experimental conditions involving the mirror. The mirror was located facing the participant at a height in the range of 15–25 cm. The distance between the participant and the mirror was 50–60 cm. Within these height and distance combinations, the spatial configuration was adjusted such that participants could gaze at the fake and actual left hands in a mirror. It should be noted that participants saw the hands from the distal direction through a mirror as shown in Fig 2(f), which is distinct from typical RHI settings.

### Tasks

In order for participants to familiarize themselves with the experimental setup, before the main tasks they wore rubber gloves and practiced tapping or rubbing a desktop surface while holding acrylic rods with a fake hand, as shown in Fig 1(a) and Fig 2(a). Participants tried to experience the illusion by tapping or rubbing (frontal-rear motion) the desk, and chose either or both of these hand movements to facilitate the induction of the illusion. During this phase, they attempted to maintain an actual right hand posture that was as that of the rubber hand. Furthermore, the fingers of the rubber hand were shaped such that the middle fingers of the actual and fake hands were at the same level and reached the desk surface simultaneously for individual participants.

Participants experienced the illusion by moving their hands for 1 min in each trial. During the tasks, participants tapped or rubbed the desktop with their actual middle fingertips for 1 min. The hand movements (rubbing stroke or tapping finger height) were limited to a few centimeters for easier manipulation and unified among all conditions for each participant. The frequency of motion was limited to approximately 1 Hz and participants were allowed to slow down their motion during the tasks to prevent fatigue and distraction. In some conditions, the acrylic rods were held such that the actual and fake hands moved synchronously. This condition causes self-generated hand motion and tactile stimuli and easily evokes the body ownership illusion [18–22]. Each experimental condition was tested in a randomized order in a single session, and two sessions were performed for individual participants. Participants were instructed to gaze at the fake hand directly in without-mirror conditions and through a mirror in with-mirror conditions.

### Experiment 1: Effect of body recognition through a mirror on the RHI

We investigated whether the experience of body ownership would be evoked when the fake hand was gazed at through a mirror. The magnitudes of the illusion with and without a mirror were compared. Three conditions were applied in this experiment, as shown in Table 1. Under conditions 1 and 2, participants held and moved the acrylic rods, and the fake and actual right hands were moved synchronously. Condition 3 was prepared as a control, and under this condition, the fake and actual hands were separated. Although the fake hand remained still on the desk, participants moved both of their hands. Participants directly gazed at the fake hand under conditions 1 and 3, and the fake hand was seen through a mirror under condition 2. Under all conditions of Experiment 1, the fake and actual hands were aligned in parallel, as shown in Fig 3(a). Each participant experienced three conditions in a randomized order in a single session, and two sessions (six trials in total) were performed for individual participants.

**Table 1.** Conditions in Experiment 1.

| No. | Condition |
| --- | --- |
| 1 | Synchronous without a mirror |
| 2 | Synchronous with a mirror |
| 3 | Asynchronous without a mirror |

**Fig 3.**
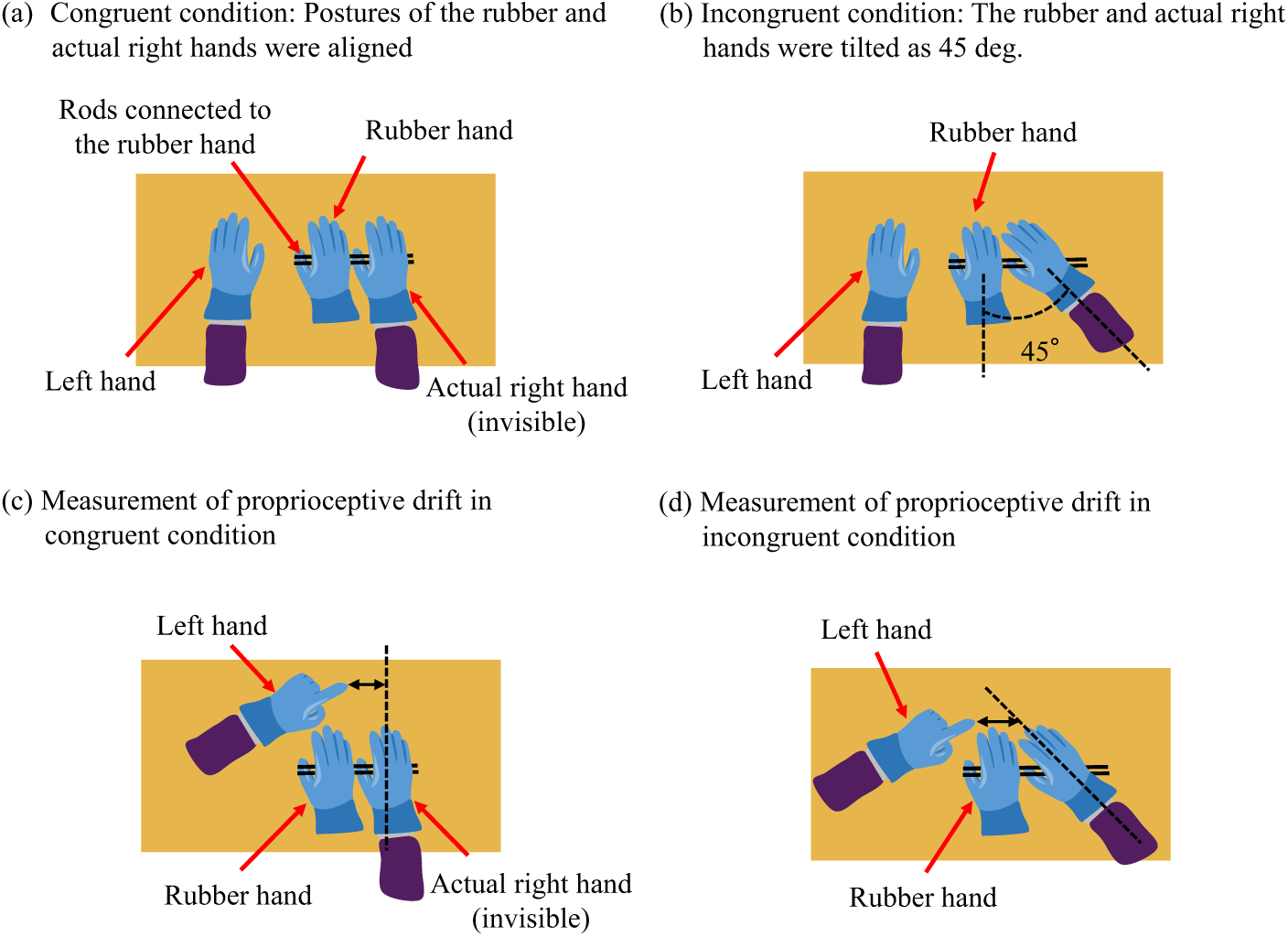
Actual and fake rubber hands. (a) The rubber hand was aligned parallel with the actual hand in Exp. 1. (b) The rubber hand was tilted 45 ° against the actual hand in Exp. 2. (c,d) The perceived finger location was measured from the longitudinal direction of the actual hand.

### Experiment 2: RHI under an incongruent hand posture with a mirror image of the fake hand

We investigated how spatial incongruency between the actual and fake hands influenced illusory body ownership. Table 2 shows the four conditions tested in Experiment 2. They included the combination of two factors: synchronization (synchronous or asynchronous) and the mirror (presence or absence). In all conditions, actual hand was tilted at 45 ° against the fake hand (Fig 3(b)). Under conditions 4 and 5, the motions of actual and fake hands were synchronized whereas they were not under conditions 6 and 7. Participants directly gazed at the fake hand under conditions 4 and 6, whereas the fake hand was seen through a mirror under conditions 5 and 7. Similar to Experiment 1, these four conditions were tested in randomized order in a single session. In total, two sessions were conducted for each individual participant.

**Table 2.** Conditions in Experiment 2.

| No. | Condition |
| --- | --- |
| 4 | Synchronous without a mirror |
| 5 | Synchronous with a mirror |
| 6 | Asynchronous without a mirror |
| 7 | Asynchronous with a mirror |

### Modified RHI questionnaire

The magnitude of an illusion is often assessed by the questionnaire proposed by Botvinick and Cohen [1]. In addition, we added questionnaire items regarding agency [23, 24] and modified some statements to fit our RHI setting. After each trial, participants responded to the six questionnaire items shown in Table 3. These items were provided in plain English. Each item was rated by using a seven-point scale (−3: I disagree with the statement, to +3: I agree with the statement). Q1 (body ownership) and Q2 (agency) concerned the illusory experience of the RHI whereas the others were prepared as controls for the body ownership statement.

**Table 3.** Questionnaire for subjective evaluation of the RHI (modified from [1])

|  |  |
| --- | --- |
| Q1 | I felt as if the seen rubber hand were my hand. |
| Q2 | It seemed that I was directly moving the seen rubber hand. |
| Q3 | It seemed as if I might have more than one right hand or arm. |
| Q4 | It felt as if my (real) hand were turning ‘rubbery’. |
| Q5 | It appeared visually as if the seen rubber hand were drifting towards right (towards my hand). |
| Q6 | The seen rubber hand began to resemble my own (real) hand, in terms of shape, size or, some other visual feature. |

### Proprioceptive drift toward the rubber hand

We measured the change of perceived finger position (proprioceptive drift) as another criterion of the RHI; however, it is known that the results do not always match with the subjective evaluation [25]. Participants pointed the position of their middle fingers of their right hands by using their index fingers of their left hands with their eyes closed. We instructed them to point 2–3 cm ahead of the fingertips of their middle fingers in a longitudinal direction to prevent them from touching the fake or actual right hands. Experimenter measured the distance between the actual and pointed positions of the middle finger in a parallel direction (Fig 3(c)). This measurement criterion was same when the actual right hand was tilted 45 ° (Fig 3(d)). PD was defined as the error of perceived finger position between the values before and after each trial. The drift toward the fake hand was considered as a positive value of PD.

## Results

We computed the mean value and standard deviation of responses toward each questionnaire item and PD of all participants by using the results of second session in each experiment.

### Results of Experiment 1

Fig 4 shows the questionnaire responses and PD in Experiment 1. We applied *t*–tests between every two conditions to verify the significant differences between them. The significance level was compensated by using the Bonferroni method.

**Fig 4.**
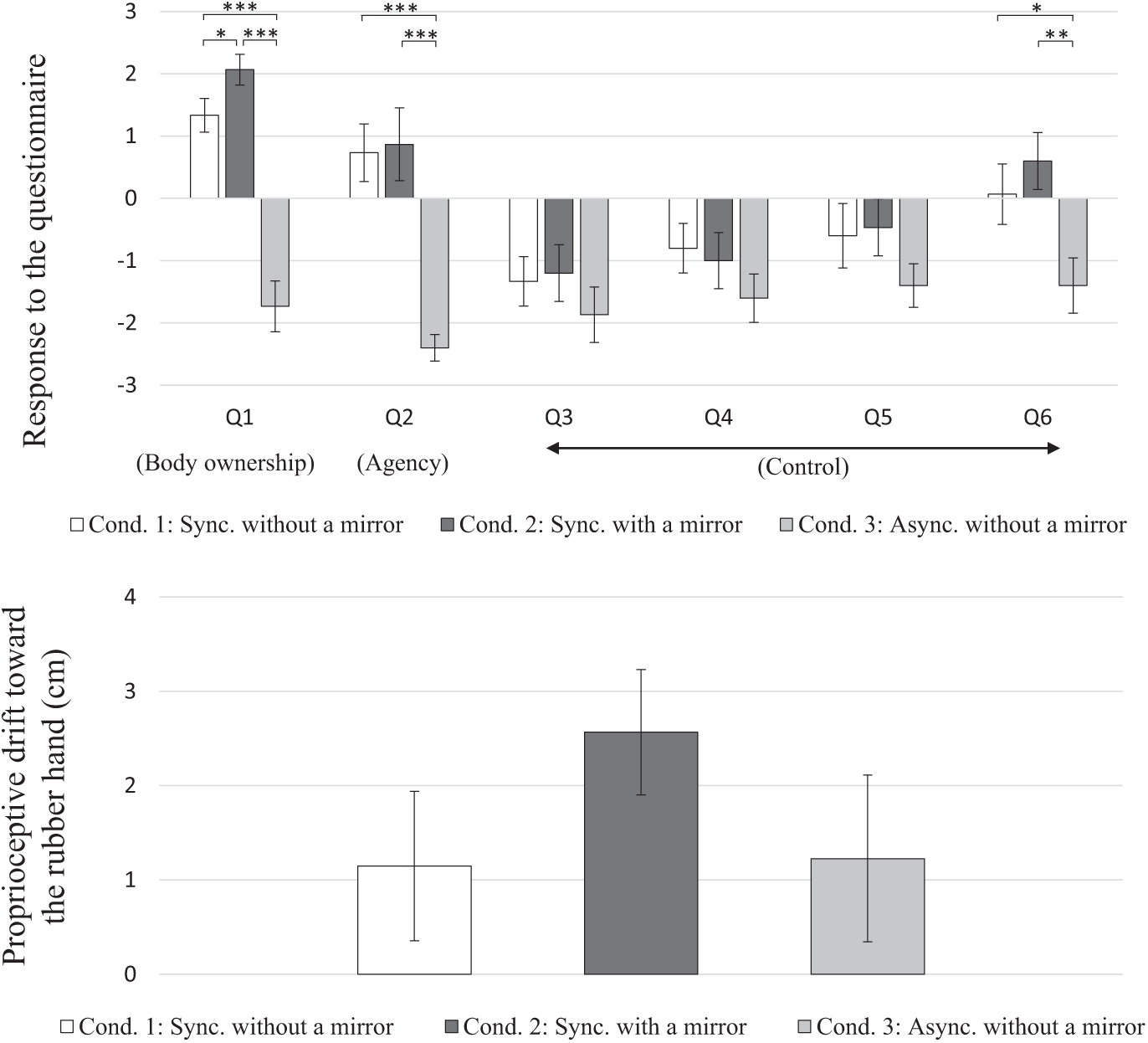
Participants’ responses to the questionnaire and proprioceptive drift toward the rubber hand in Exp. 1. Means and standard errors among the participants. Asterisks *, **, and *** indicate *p <* 0.05, *p <* 0.01, and *p <* 0.001, respectively using the Bonferroni correction. Positive values of PD show that the position of the perceived hand was close to that of the fake hand.

#### Questionnaire scores

The scores of Q1 (body ownership) at the condition with a mirror were significantly greater than those without a mirror under synchronous conditions (Cond. 1 vs Cond. 2, *t*(14) = 2.75, *p* = 0.047 < 0.05). The synchronization significantly increased the score of Q1 under gazing at the fake hand directly (Cond. 1 vs Cond. 3, *t*(14) = 6.78, *p* < 0.001). Significant differences were also confirmed between ‘mirror–synchronous’ condition and ‘direct view–asynchronous’ condition (Cond. 2 vs Cond. 3, *t*(14) = 7.21, *p* < 0.001).

In terms of Q2 (agency), the effect of the mirror was not significant under synchronous conditions (Cond. 1 vs Cond. 2, *t*(14) = 0.254, *p* > 0.05, n.s.), whereas the scores were positive regardless of the presence of mirror. Same as the result of Q1, the effect of synchronization was confirmed under directly gazing at the fake hand (Cond. 2 vs Cond. 3, *t*(14) = 5.14, *p <* 0.001), and significant differences were confirmed between the ‘mirror–synchronous’ condition and ‘direct view–asynchronous’ condition (Cond. 1 vs Cond. 3, *t*(14) = 5.52, *p <* 0.001).

These results indicate that illusory body ownership was experienced when the mirror was adopted. Body ownership was strongly induced by the use of a mirror and agency was induced regardless of the presence of a mirror under synchronous conditions.

The scores of Q3–5 (i.e., control items) were negative under all conditions. However, in terms of Q6, the scores under synchronous conditions (Cond. 1 and Cond. 2) were greater than those under asynchronous conditions (Cond. 3).

#### Proprioceptive drift

The mean values of PD were positive for all conditions, and that for the ‘mirror–synchronous condition’ (Cond. 2) was the greatest. However, significant differences were not confirmed between every pair of conditions.

### Results of Experiment 2

Fig 5 shows the questionnaire responses and PD in Experiment 2. We applied two-way analysis of variance (ANOVA) to the results using the presence/absence of the mirror and synchronization as the factors.

**Fig 5.**
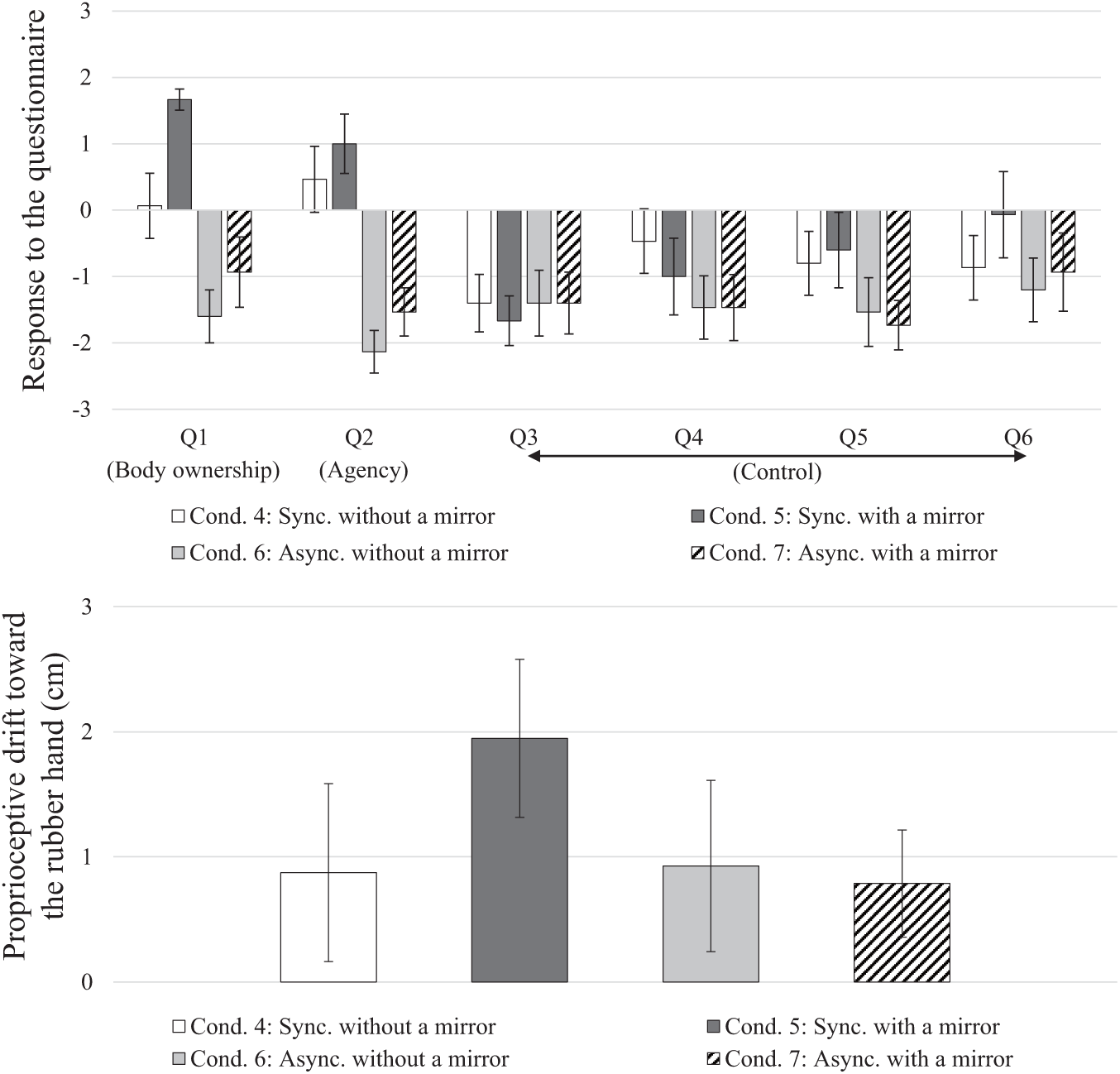
Participants’ responses to the questionnaire and proprioceptive drift toward the rubber hand in Exp. 2. Means and standard errors among the participants. Positive values of PD show that the position of the perceived hand was close to that of the fake hand.

#### Questionnaire scores

The scores of Q1 (body ownership) were significantly greater when using a mirror (*F*_1,59_ = 7.25, *p* < 0.01), or when the hand motions were synchronized (*F*_1,68_ = 25.7, *p* < 0.001), whereas an interaction effect between them was not detected (*F*_1,59_ = 1.23, *p* > 0.05, n.s.).

The scores of Q2 (agency) were significantly greater under synchronous conditions (*F*_1,59_ = 38.64, *p* < 0.001), whereas the effect of the mirror (*F*_1,59_ = 1.88, *p* > 0.05, n.s.) and their interaction effect (*F*_1,59_ = 0.01, *p* > 0.05, n.s.) were not detected.

For incongruent conditions, body ownership was strongly induced when a mirror was adopted and agency was induced regardless of the presence of mirror. The scores of control items were negative for all conditions.

#### Proprioceptive drift

The mean values of PD were positive under all conditions and that of the ‘mirror–synchronous condition’ (Cond. 5) was the greatest. The effects of the mirror (*F*_1,59_ = 0.56, *p* > 0.05, n.s.), the synchronization (*F*_1,59_ = 0.79, *p* > 0.05, n.s.), and their interaction (*F*_1,59_ = 0.94, *p* > 0.05, n.s.) were not significant. PD was not significantly influenced by either the presence/absence of the mirror or by the synchronization.

## Discussion

In Experiment 1, illusory body ownership was experienced by participants when seeing the fake hand in a mirror. Body ownership was significantly strengthened when a mirror was adopted whereas agency was induced regardless of the presence of mirror under synchronous conditions with a congruent hand posture. No significant differences were confirmed from the magnitude of PD in terms of the presence of the mirror and synchronization. These results are supported by the study of Jenkinson and Preston [12], which investigated the mirror RHI. Other related studies have reported body ownership through a mirror at the same extent [10] or weaker [11] as compared to the condition without a mirror. However, these studies investigated passive RHI, in which the experimental conditions differed from our study and that of study of Jenkinson and Preston [12] in terms of involvements of hand movements.

In Experiment 2, ownership over the fake hand was observed even when the fake and actual hands were spatially incongruent in terms of the hand angle. For incongruent conditions when directly gazing at the fake hand (Cond. 4), body ownership was weak whereas agency was induced. Postural incongruency tends to weaken body ownership but does not affect agency in active RHI [23, 24]. However, when using a mirror (Cond. 5), stronger body ownership and the same extent of agency were induced compared with condition without a mirror. This indicates that even under incongruent conditions, not only agency but also body ownership can be induced by using a mirror.

The effect of the mirror on the illusion was detected regardless of synchronization under incongruent conditions. Many studies report that the illusion does not occur under asynchronous conditions. Although some readers may doubt the effect of suggestibility, the subjective scores against asynchronous conditions are all negative. Thus, neither body ownership nor agency would have been induced, or the self-body recognition was obscured by a mirror and this might have facilitated the illusion even for ‘mirror–asynchronous’ condition. Humans are likely to misunderstand their body motion seen through a mirror [26], and transformation into a third-person view decreases the precision of visual and motor perception [27]. Many studies have discussed the differences between first-person and third-person views, and suggested their effects on the self-body recognition [10, 12, 28].

The enhancement of the illusion by a mirror could be explained by two possible reasons if self-body recognition is obscured by a mirror. First, the mirror decreases the reliability and precision of visual cues and masks the inconsistency of perceived signals. Provided that the perceived hand is determined by a multisensory integration processes of sensory cues [29–31], noisy or imprecise images in a mirror may alter the weightings of unisensory signals for multisensory integration [12]. For instance, in a visual-haptic integration task, a decrease in the reliability of visual cues led to the relative increase in the contribution of haptic cues [32–34]. In particular, body ownership is thought to be induced in the process of inferring the environment from perceived signals [35]. Therefore, illusory body ownership would be influenced by the change of weightings of multisensory cues. In our experimental settings, visual, tactile, and proprioceptive cues were involved, and the use of a mirror degraded reliability regarding the visually provided hand posture. The weightings of sensory cues would be different between conditions with and without a mirror. Such an effect of less reliable images in a mirror was pointed out in [11], in which even laterality was masked by using mirror images of a fake hand. In other words, in classical RHI settings where the fake hand is gazed at directly, because of the greater contribution of visual cues, visible incongruency may cause the visual sense to fail to capture other cues and prevent the occurrence of the illusion.

The second reason is that visual cues are weakened by a mirror and this accelerates multisensory integration. The body-ownership illusion is considered to be caused by visuo-tactile integration, and a greater degree of integration would produce a more intense illusion. Sensory integration under RHI settings has been demonstrated in terms of spatial and temporal accordance [3–5]. In addition, multisensory integration is enhanced when the stimuli to individual sensory channels are weak [36, 37]. These three aspects are known as the spatial rule, temporal rule, and inverse effectiveness [38], which was studied neurophysiologically from the perspective of the activity of superior colliculus cells [36] and inter-trial phase coherency of scalp EEG (electroencephalogram) during RHI tasks [39, 40]. In our experiment, the use of a mirror might have weakened the visual stimuli and could be related to the principle of inverse effectiveness, and multisensory integration might have been enhanced in comparison with using the settings without a mirror.

## Conclusions

This study investigated the role of gazing at a fake hand in a mirror under RHI conditions. The subjective experience was assessed based on a questionnaire. The illusory experiences of body ownership and agency were evoked when the fake hand in a mirror was gazed at (Experiment 1). The magnitudes of subjective experiences were higher (body ownership) or equally high (agency) compared with the condition where the fake hand was directly seen without a mirror. Furthermore, using the mirror, the illusory experience was reported even when the actual and fake hands were placed at incongruent postures (Experiment 2). Although the underlying mechanism of these observations is unknown, gazing at a fake hand in a mirror offers a promising approach for robustly inducing intense RHI experiences.

## Acknowledgments

We thank Ariei Oishi and Atsuko Tamada for their assistance in conducting the experiments.

